# Arabidopsis RNA Polymerase IV generates 21-22 nucleotide small RNAs that can participate in RNA-directed DNA methylation and may regulate genes

**DOI:** 10.1101/832634

**Authors:** Kaushik Panda, Andrea D. McCue, R. Keith Slotkin

## Abstract

The plant-specific RNA Polymerase IV (Pol IV) transcribes heterochromatic regions, including many transposable elements, with the well-described role of generating 24 nucleotide (nt) small interfering RNAs (siRNAs). These siRNAs target DNA methylation back to transposable elements to reinforce the boundary between heterochromatin and euchromatin. In the male gametophytic phase of the plant life cycle, pollen, Pol IV switches to generating primarily 21-22 nt siRNAs, but the biogenesis and function of these siRNAs has been enigmatic. In contrast to being pollen-specific, we identified that Pol IV generates these 21-22 nt siRNAs in sporophytic tissues, likely from the same transcripts that are processed into the more abundant 24 nt siRNAs. The 21-22 nt forms are specifically generated by the combined activities of DICER proteins DCL2/DCL4 and can participate in RNA-directed DNA methylation. These 21-22 nt siRNAs are also loaded into ARGONAUTE1, which is known to function in post-transcriptional regulation. Like other plant siRNAs and microRNAs incorporated into AGO1, we find a signature of genic mRNA cleavage at the predicted target site of these siRNAs, suggesting that Pol IV-generated 21-22 nt siRNAs may function to regulate gene transcript abundance. Our data provides support for the existing model that in pollen Pol IV functions in gene regulation.

## Background

Transposable elements (TEs) are mobile DNA fragments that cause mutations by inserting into genes and creating chromosomal breaks. To repress their mobility, and therefore limit the number of new mutations, eukaryotes target TE activity at the transcriptional, post-transcriptional and translational levels (reviewed in [1]). A major regulatory mechanism used to repress TEs are small RNAs, which target TE mRNAs for degradation, inhibit the translation of TE protein, and can guide *de novo* chromatin modification of TE loci, resulting in transcriptional silencing. In flowering plants, TE small interfering RNAs (siRNAs) are well-studied and fall into two major categories: 21-22 nucleotide (nt) siRNAs generated by RNA Polymerase II (Pol II), and 24 nt siRNAs generated by the plant specific RNA Polymerase IV (Pol IV)(reviewed in [2]).

Early in plant evolution, the protein subunits of the Pol II holoenzyme duplicated and subfunctionalized into two additional RNA polymerase complexes, Pol IV and Pol V [3]. The biological function of Pol IV is to transcribe heterochromatic regions of plant genomes into non-polyadenylated transcripts that are created for the sole purpose of siRNA generation [4]. Pol IV is guided to heterochromatic target regions of the genome by the mark of histone H3 lysine 9 dimethylation (H3K9me2) [5], and creates short 26-45 nt transcripts that are converted into double-stranded RNA via the RNA-DEPENDENT RNA POLYMERASE 2 protein (RDR2) [6,7]. This double-stranded RNA is then cleaved by DICER-LIKE 3 (DCL3) into predominantly 23-24 nt siRNAs [8]. Pol IV-derived 24 nt siRNAs are incorporated into the ARGONAUTE4 (AGO4) and AGO6 proteins, to guide AGO function in RNA-directed DNA methylation (RdDM) of the target locus [9]. Therefore, Pol IV’s overall function in plant biology is to generate the siRNAs necessary to reinforce heterochromatic marks and maintain euchromatin/heterochromatin boundaries [10]. A secondary role is to generate an siRNA defense against any new or active TEs that share sequence homology [11].

Pollen is the male gametophytic generation of flowering plants, and contains two sperm cell gametes encapsulated in a larger vegetative cell, which directs the delivery of the sperm cells upon pollination. Pollen has a known broad activation of TE expression resulting in steady-state TE mRNAs, which occurs from the nucleus of the pollen vegetative cell [12,13]. This TE activation occurs simultaneously with abundant TE 21-22 nt siRNA production in the pollen grain [12,14,15]. Other cases of TE transcriptional activation in the sporophytic plant body, for example in mutants of the TE master chromatin modifying gene DDM1, are also associated with 21-22 nt siRNA production from Pol II-derived TE mRNAs [16]. These siRNAs were termed *epigenetically-activated siRNAs* (*easiRNAs*) because they appear only when TEs lose transcriptional repression and produce Pol II-derived mRNAs [15,17]. It was therefore assumed that in pollen the reactivated TE-generated Pol II mRNAs were the source of pollen easiRNAs [12]. However, a recent publication demonstrated that in the pollen of wild-type reference ecotype *Columbia* plants (wt Col), *pol IV* mutants fail to generate 21-22 nt TE siRNAs [15]. This suggested a key role of Pol IV beyond the known production of 24 nt siRNAs.

We aimed to determine whether pollen 21-22 nt easiRNAs are actually produced from Pol IV transcripts, or alternatively whether Pol IV is necessary to trigger siRNA production from Pol II transcripts. We found that pollen TE easiRNA production is a product of Pol IV transcription, and this activity of Pol IV is not specific to pollen. We find that in the absence of the more abundant 24 nt siRNAs, Pol IV-derived 21-22 nt siRNAs can participate in RdDM. Like other 21-22 nt siRNAs generated from Pol II, Pol IV 21-22 nt siRNAs are incorporated into AGO1, which is the main effector protein of post-transcriptional silencing. Our data suggests that like other siRNAs and microRNAs incorporated into AGO1, Pol IV-dependent 21-22 nt siRNAs may participate in the post-transcriptional targeting of genic mRNAs.

## Results

### Pol IV is required for production of TE 21-22 nt siRNAs

Arabidopsis easiRNAs were discovered in gametophytic pollen and found to be primarily 21-22 nt in length. In contrast, heterochromatic siRNAs are produced during sporophytic stages and are primarily 24 nt in length. To compare the change in siRNA size distribution during development, we analyzed small RNAs sequenced from wt Col seedling [18], inflorescence (this study) and pollen [15]. We used diploidized 2n wt Col pollen (derived from 4n wt Col) as a second pollen replicate (see Methods). We confirmed that compared to seedling and inflorescence, there is a sharp increase in relative amounts of TE 21-22 nt siRNAs and a corresponding decrease of TE 24 nt siRNAs in pollen (Figure 1A). However, we found that the shift in small RNA size relative accumulation of Figure 1A was primarily due to a sharp decrease in TE 24 nt siRNA production in pollen, and not an increase in TE 21-22 nt siRNA abundance (Figure 1B).

**Figure 1.**
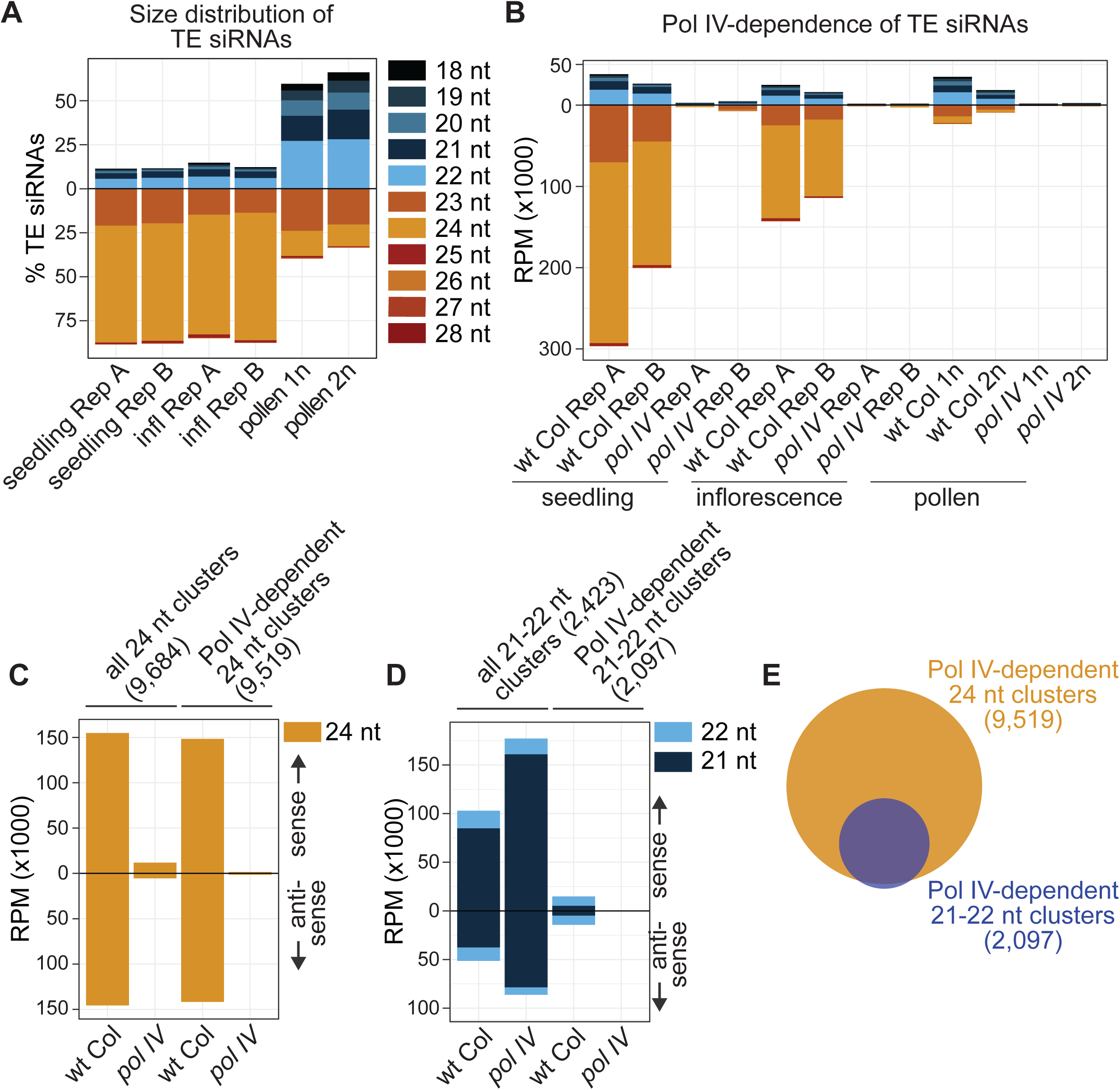
Pol IV-dependent 21-22 nt siRNAs. A. Percent of TE siRNAs in different size classes in wild-type Columbia (wt Col) seedlings, inflorescence and pollen tissue. Rep A and Rep B are distinct biological replicates, 1n is normal haploid pollen, and 2n is diploid pollen derived from a 4n parent [15]. B. Reads per million (RPM) counts of TE siRNAs in wt Col and *pol IV* mutant plants. C. 24 nt siRNA accumulation in inflorescence from all identified 24 nt siRNA-producing clusters (left) or clusters specifically identified as dependent on Pol IV (right). Above zero are sense siRNAs, and below are antisense. D. 21-22 nt siRNA accumulation in inflorescence from all identified 21-22 nt siRNA-producing clusters (left) or clusters identified that are dependent on Pol IV (right). E. Overlap between the regions of the genome where Pol IV produces 24 nt vs. 21-22 nt siRNAs.

The recently reported Pol IV-dependence of 21-22 nt siRNAs in pollen [15] was unexpected, since Pol IV had previously only been shown to generate 23-24 nt siRNAs [19](reviewed in [20]). Therefore, we aimed to determine if Pol IV-dependent TE 21-22 nt siRNA production is specific to pollen or occurs in non-gametophytic tissues. We confirmed that both 21-22 and 24 nt TE siRNAs are dependent on Pol IV in pollen (Figure 1B) [15]. We also identified that in the TE-silent sporophytic seedling and inflorescence tissue, Pol IV is responsible for the accumulation of TE 21-22 nt siRNAs (Figure 1B). For comparison across different tissue types and sequencing libraries, we normalized the TE siRNA counts by total sequenced small RNAs that match the Arabidopsis genome (Figure 1A-B). To confirm that these observations were not biased due to our specific normalization method, we alternatively normalized TE siRNAs using Pol IV-independent miRNA counts (Supplementary Figure 1). We find that the reduction in accumulation of TE siRNAs in *pol IV* mutants is consistent regardless of normalization method. We conclude that Pol IV-dependent TE 21-22 nt siRNAs are not specific to pollen, and a similar mechanism of 21-22 nt siRNA production exists during sporophytic stages.

To take an unbiased approach to investigate all small RNAs (sRNAs) beyond annotated TEs, we identified clusters of 24 nt sRNAs and 21-22 nt sRNAs in wt Col inflorescence (Figure 1C-D). Expectedly, almost all 24 nt sRNAs are lost from the 24 nt clusters in a *pol IV* mutant (Figure 1C). In contrast, global levels of 21-22 nt sRNAs increase in *pol IV* mutants (Figure 1D), which has been previously reported [21]. A majority of these 21-22 nt sRNAs are miRNAs and/or miRNA-induced (tasiRNAs) which are not dependent on Pol IV production. This increased overall level of 21-22 nt sRNAs has likely obscured the fact that Pol IV-dependent 21-22 nt sRNA regions of the genome do exist in wt Col inflorescence (Figure 1D), accounting for why they were not discovered earlier. In addition, >99% of Pol IV-dependent 21-22 nt clusters overlap with 24 nt clusters (Figure 1E). We conclude that there is a genome-wide population of Pol IV-dependent 21-22 nt siRNAs, so far uninvestigated, which are generated from a subset of loci that also produce Pol IV-dependent 24 nt siRNAs.

### Pol IV-dependent 21-22 nt siRNAs are produced from Pol IV transcripts

It is well established that 24 nt siRNAs are produced from Pol IV transcripts (reviewed in [20]). We aimed to determine whether 21-22 nt siRNAs are also produced from Pol IV transcripts, or are instead produced from Pol II transcripts but somehow dependent on Pol IV. As shown in Figure 1E, almost all 21-22 nt clusters overlap with 24 nt siRNAs clusters, suggesting that 21-22 nt siRNAs are produced from a subset of 24 nt clusters and therefore likely from Pol IV transcripts. To investigate the extent of overlap between the two clusters, we positioned each 21-22 nt cluster relative to its corresponding overlapping aligned 24 nt siRNA cluster (Figure 2A). We found that most of the 21-22 nt clusters (92%) aligned within the boundaries of 24 nt clusters. Upon investigation of the remaining (8%) 21-22 nt clusters, we found that these loci also shadow 24 nt siRNA loci, but were falsely classified as extending beyond 24 nt clusters due to bioinformatic artefacts of cluster identification. Therefore, we failed to identify any locus producing Pol IV-dependent 21-22 nt siRNAs that does not also produce 24 nt siRNAs. This observation strongly suggests that Pol IV transcripts that feed into the 24 nt siRNA pathway also produce 21-22 nt siRNAs.

**Figure 2.**
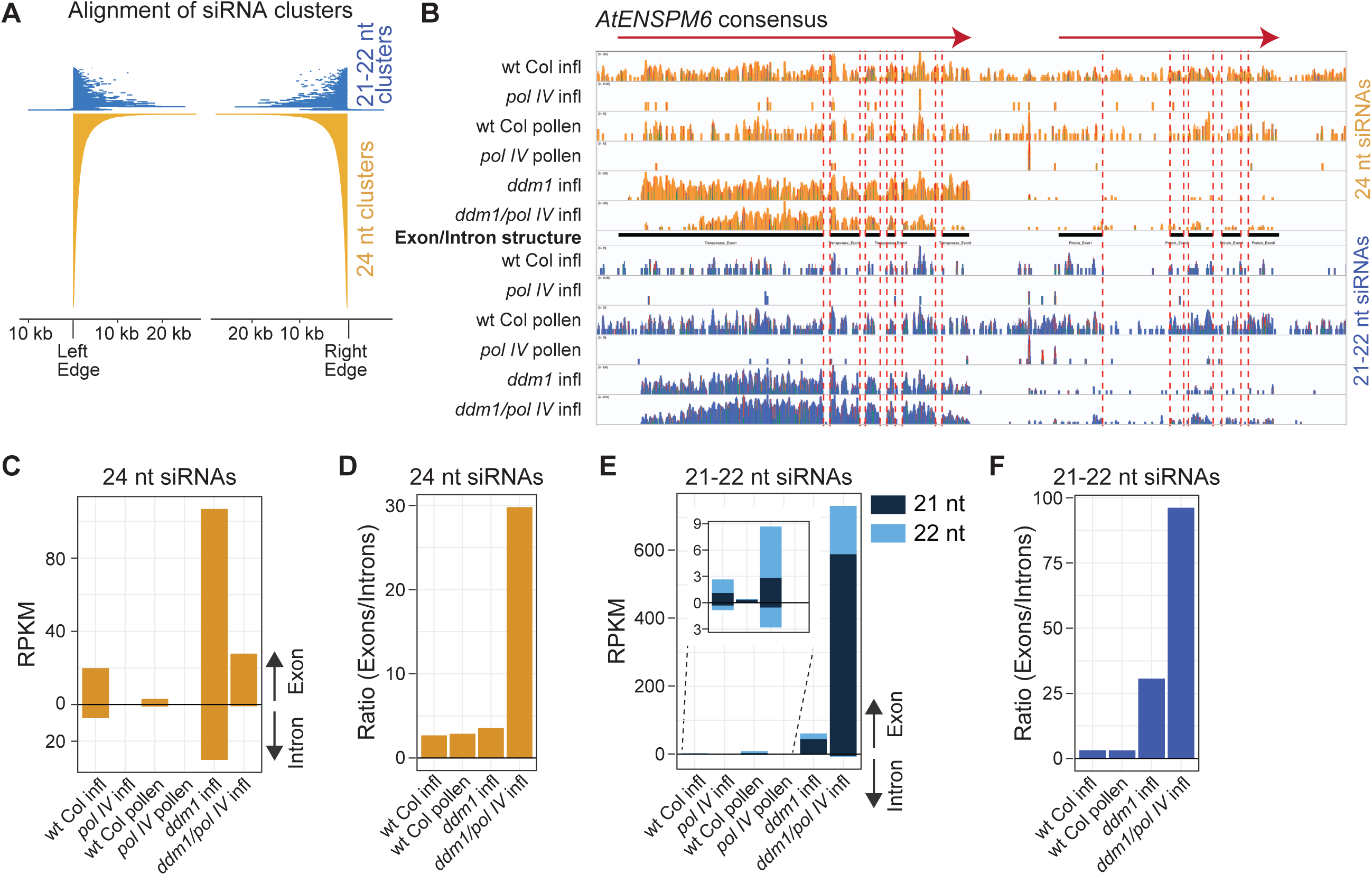
Pol IV-dependent 21-22 nt siRNAs are produced from Pol IV transcripts. A. Alignment of the 21-22 nt siRNA clusters to the ordered alignment of the 24 nt siRNA clusters. Very few 21-22 nt clusters extend beyond the boundaries of the 24 nt clusters, and inspection of these found no confirmable region of the genome that generates Pol IV-dependent 21-22 nt siRNAs but not 24 nt siRNAs in inflorescence tissue. B. Alignment of siRNAs from various genotypes and tissues to the *AtENSPM6* TE consensus sequence, which encodes two transcripts (red arrows). Black bars indicate protein-coding exons. Red dashed lines indicate transcript intron boundaries. Inflorescence is abbreviated as infl. C. Read counts of intron- and exon-aligning 24 nt siRNAs for *AtENSPM6*. D. Ratio of exon to intron small RNA abundance for 24 nt siRNAs. E. Read counts of intron- and exon-aligning 21-22 nt siRNAs for *AtENSPM6*. Inset shows the low-level accumulation when TEs are not transcriptionally activated in the *ddm1* mutant. F. Ratio of exon to intron small RNA abundance for 21-22 nt siRNAs.

To further investigate the origin of Pol IV-dependent siRNAs, we used the exon-intron structure of transcripts that produce siRNAs. Pol II-transcribed RNAs are efficiently co-transcriptionally spliced and are therefore cleaved into siRNAs matching only exons (Figure 2B-F), but the exon-intron distribution of Pol IV siRNAs has not been investigated. We focused on the consensus sequence of the *AtENSPM6* family of TEs and its annotated exon-intron structure in the GIRI Repbase [22]. We aligned both 21-22 and 24 nt siRNAs from six key plant lines to this consensus sequence (Figure 2B) and counted the abundance from exons and introns (Figure 2C,E). We also calculated the ratio of exonic/intronic siRNAs (Figure 2D,F) and aimed to use the relative bias in exonic/intronic ratio as a signature for determining the polymerase origin of siRNAs.

We found that when TEs are transcribed by Pol II while Pol IV is not present (*ddm1 pol IV* double mutant), siRNAs have a high exon/intron bias for both 24 nt and 21-22 nt siRNAs (Figure 2B-F). In this double mutant, when Pol II is the only polymerase generating TE siRNAs, the level of exon reads outweighs intron reads 30X for 24 nt siRNAs, and 95X for 21-22 nt siRNAs (Figure 2D,F). When a functional Pol IV protein is present, in the *ddm1* single mutant (both Pol IV and Pol II active), the bias of exon/intron siRNAs is severely reduced suggesting that Pol IV-derived siRNAs are produced from intronic regions as well (Figure 2C-F). This conclusion is supported by 24 nt siRNA production in TE-silent wt Col inflorescence, in which TE siRNAs are known to be produced from Pol IV transcripts (Pol IV active, but Pol II inactive at TEs), and therefore have a low exon/intron ratio bias in wt Col inflorescence (Figure 2C,D). Using these observations as controls, we investigated the exon/intron bias of 21-22 nt siRNAs in wt Col. We found that in inflorescence and pollen the Pol IV-dependent 21-22 nt siRNAs have a low bias of exon/intron siRNAs (Figure 2E,F), therefore demonstrating that unspliced and likely Pol IV transcripts produce 21-22 nt siRNAs in TE-silent wt Col inflorescence and in TE-active pollen. Together with the observation that 21-22 nt siRNAs are completely lost in *pol IV* (Figure 2C,E), we conclude that Pol IV (and not Pol II) produce the Pol IV-dependent 21-22 nt siRNAs.

### Pol IV-dependent 21-22 nt siRNAs are produced by DCL2 and DCL4

To address whether the Pol IV derived 21-22 nt siRNAs are non-specific degradation products produced from Pol IV transcripts or full-length 24 nt siRNAs, we compared siRNA accumulation in wt Col and DCL protein family mutants. As expected, we observed the complete loss of 24 nt siRNAs in the *dcl3* mutant (Figure 3A) [8]. DCL family proteins have known redundancies [23], so when DCL3 is absent, DCL2 and DCL4 substitute and process Pol IV transcripts into 21-22 nt siRNAs (Figure 3A,B), confirming that DCL2 and DCL4 have the ability to process Pol IV transcripts [8]. When specifically focused on Pol IV 21-22 nt clusters (Figure 3B), we observe a class of Pol IV-dependent 21-22 nt siRNAs in wt Col that are dependent on DCL2 and DCL4 for their production (inset, Figure 3B). This demonstrates that even in wt Col sporophytic tissue, Pol IV generates 21-22 nt siRNAs that are not random degradation products of Pol IV transcripts or full-length 24 nt siRNAs, but rather are specific cleavage products of DCL2 and DCL4. We conclude that Pol IV transcripts are acted upon by DCL2, DCL3 and DCL4, with a strong bias towards DCL3 and production of 24 nt siRNAs in sporophytic tissues.

**Figure 3.**
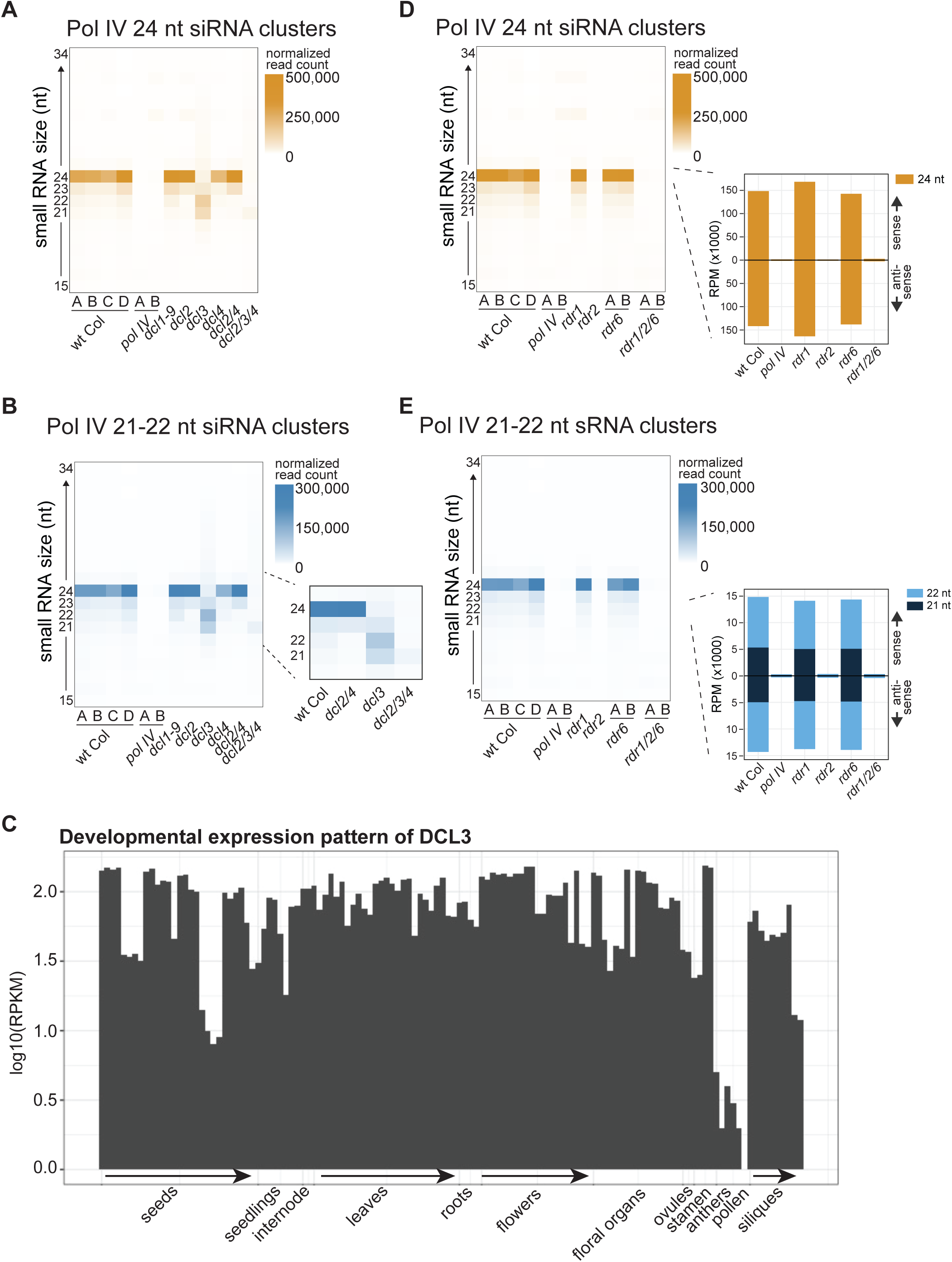
Pathway for Pol IV-dependent siRNA production. Heatmap of read counts of 15-34 nt small RNAs in inflorescence from Pol IV-dependent clusters in various mutant combinations. 24 nt siRNAs clusters are used in (A,D) and 21-22 nt siRNA clusters are used in (B,E). (A-B) uses DCL family mutants. *dcl2/3/4* refers to the *dcl2/dcl3/dcl4* triple mutant. (D-E) uses RDR family mutants, and *rdr1/2/6* refers to the *rdr1/rdr2/rdr6* triple mutant. Insets of D and E show accumulation of 24 nt and 21-22 nt siRNAs respectively with abundance averaged across replicates. C) Developmental expression pattern of DCL3 protein according to [41] and analyzed in the eFP-RNA-Seq Browser [42].

The WT pollen siRNA profile of relatively equal amounts of 21, 22 and 24 nt Pol IV-derived siRNAs (Figure 1B) could be produced by a partial absence or dysfunction of the DCL3 protein, whereby more Pol IV transcripts are processed by DCL2 and DCL4. We examined DCL3 expression in pollen and found a lack of expression specifically in this tissue (Figure 3C). This suggests that DCL3 protein may be simply lacking or reduced in WT pollen, however, mRNAs of many siRNA-generating proteins are reduced or absent in WT pollen (including the largest subunit of Pol IV itself (NRPD1), Supplemental Figure 3). There is a poor correlation between steady-state mRNA levels and protein abundance (reviewed in [24]), and that further analysis is needed to confirm if reduction in DCL3 activity is responsible for the pollen-specific TE siRNA size distribution.

We next aimed to identify a DCL mutant combination that removes all siRNA production from Pol IV transcripts. The overall abundance of siRNAs is reduced in *dcl2/3/4* triple mutants, however 21 nt siRNAs are still detected (Figure 3A,B). To investigate the source of 21 nt siRNAs in *dcl2/3/4*, which was assumed to be DCL1, we interrogated a seedling tissue dataset that includes mutations in all four DCL family proteins (*dcl1/2/3/4*) [18]. Using this dataset, we first confirmed that the Pol IV-dependent siRNA producing clusters identified in inflorescence, also produce Pol IV-dependent siRNAs in seedling (Supplementary Figure 2). Second, the loss of 24 nt siRNAs and increase in relative abundance of 21 nt siRNAs in *dcl2/3/4* at the Pol IV-dependent siRNA clusters is also observed in seedlings. We find that these 21 nt siRNAs are produced from Pol IV transcripts, since these siRNAs are lost in *pol IV dcl2/3/4* (compared to *dcl2/3/4*). However, we found that all siRNAs are not lost in *dcl1/2/3/4* quadruple mutants (Supplementary Figure 2A-B), which therefore must be due to the DICER-independent pathway of Pol IV small RNA production [18].

In Figure 1 we found a relatively equal distribution of Pol IV-dependent 21-22 nt siRNAs between sense and antisense strands (Figure 1D), suggesting that like 24 nt siRNAs (Figure 1C), 21-22 nt siRNAs are produced from double-stranded Pol IV transcripts. To further elucidate the pathway of siRNA biogenesis, we investigated the production of siRNAs in plants mutated for RDR family proteins. We found that both 21-22 and 24 nt siRNAs are dependent on RDR2, and are unperturbed in either *rdr1* or *rdr6* single mutants (Figure 3D-E). Therefore, Pol IV-derived 21-22 nt siRNA production is distinct from Pol II-derived TE siRNA production (in *ddm1* mutants), which requires RDR6 [16]. We conclude that in sporophytic tissues, Pol IV / RDR2 generates double-stranded TE transcripts that are primarily cleaved into 24 nt siRNAs by DCL3, but are also cleaved by DCL2/DCL4 into low levels of 21-22 nt siRNAs.

### Pol IV 21-22 nt siRNAs can target RNA-directed DNA methylation

Pol IV-derived 24 nt siRNAs have well-established roles in guiding RdDM [9]. To determine if Pol IV-derived 21-22 nt siRNAs can function in RdDM, we used MethylC-seq to assay genome-wide DNA methylation in a series of DCL family single, double and triple mutants. We identified differentially methylated regions (DMRs) in *pol IV* mutant plants and aligned CHH context DNA methylation (H=A,C or T) at their edge (Figure 4A). Asymmetric CHH methylation, particularly at Pol IV DMRs, is a hallmark of the RdDM pathway [25]. Importantly, the methylation level of the *dcl3* single mutant is not as low as the *pol IV* mutant (Figure 4A), demonstrating that Pol IV-dependent methylation can function through other DCL proteins. In addition, the methylation level in *dcl3* is not as low as in as the *dcl2/3/4* triple mutant, demonstrating that specifically DCL2 and DCL4 have a function in targeting DNA methylation (Figure 4A). SiRNAs from these same regions show the increased abundance of 21-22 nt siRNAs in the *dcl3* mutant (Figure 4B). These siRNAs must participate in RdDM, as they are lost in *dcl2/3/4* (Figure 4B), resulting in reduced methylation (Figure 4A). Figure 4C demonstrates that the loss of methylation in Figure 4A is not the product of just a few loci. Together, these data demonstrate that in the absence of DCL3 and 24 nt siRNA production, Pol IV-generated 21-22 nt siRNAs can participate in RdDM.

**Figure 4.**
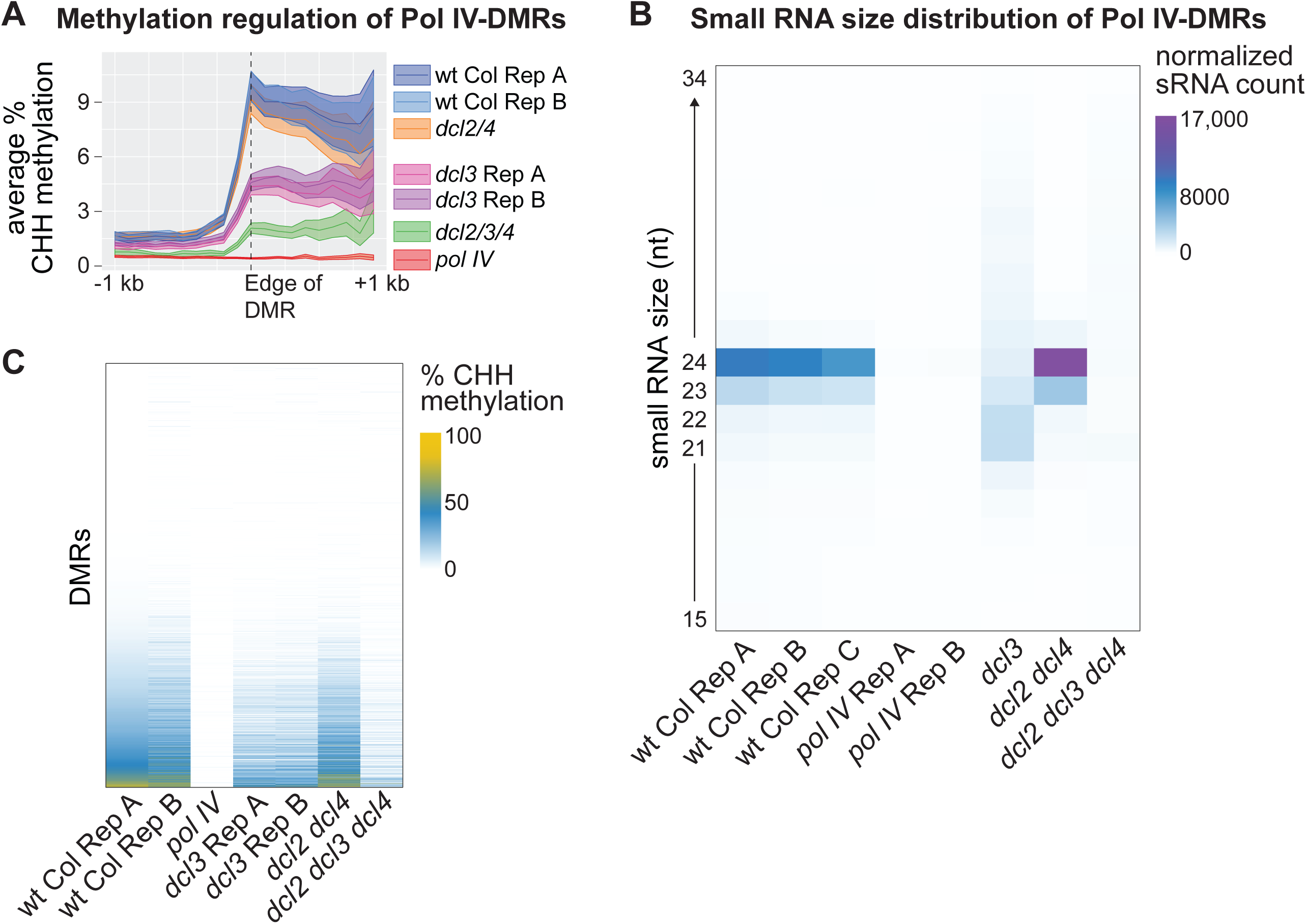
Pol IV-dependent 21-22nt siRNAs can cause DNA methylation. A. Metaplot of average percent CHH context methylation of Pol IV-DMRs in wt Col and different mutant combinations. B. Small RNA size distribution of the Pol IV-DMRs shown in A. C. Heatmap of CHH DNA methylation at Pol IV-DMRs where each row represents an individual DMR. DMRs are sorted by their methylation level in WT Col Rep A.

### Pol IV 21-22 nt siRNAs may target gene transcripts

Given the size of Pol IV-dependent 21-22 nt siRNAs, and the known function of tasiRNAs, microRNAs, and TE siRNAs of this size to act in post-transcriptional gene regulation (PTGS) of genic mRNAs, we wondered if Pol IV-dependent 21-22 nt siRNAs played a similar role. If Pol IV-derived 21-22 nt siRNAs target genic transcripts for post-transcriptional regulation, then there are four predictions to be tested.

First, these Pol IV-derived 21-22 nt siRNAs would be incorporated into the genic mRNA-regulating AGO protein, AGO1. Pol II-derived 21-22 nt siRNAs are known to cleave mRNA transcripts of genes *in trans* through the effector protein AGO1 [26]. We investigated whether Pol IV-derived siRNAs are incorporated into AGO1. We sequenced siRNAs from AGO1 immuno-precipitations (IPs) in both wt Col and *pol IV* mutants. As a control, we also sequenced siRNAs from no-antibody controls (mock IPs) in both wt Col and *pol IV.* Positive and negative controls that ensure that our IP sRNA-seq experiment worked as expected are shown in Supplemental Figure 4. We found that like Pol II-derived 21-22 nt microRNAs, tasiRNAs and some 21-22 nt TE siRNAs [27], Pol IV-derived 21-22 nt siRNAs are enriched in AGO1 (Figure 5A). As expected, these AGO1-incorporated 21-22 nt siRNAs are completely lost in *pol IV* mutants. As a control, we confirmed that Pol IV-derived 24 nt siRNAs are not strongly enriched in AGO1 (Figure 5B). AGO1 incorporation suggests that Pol IV siRNAs could act to target the known activity of AGO1 for mRNA transcript cleavage and translational inhibition.

**Figure 5.**
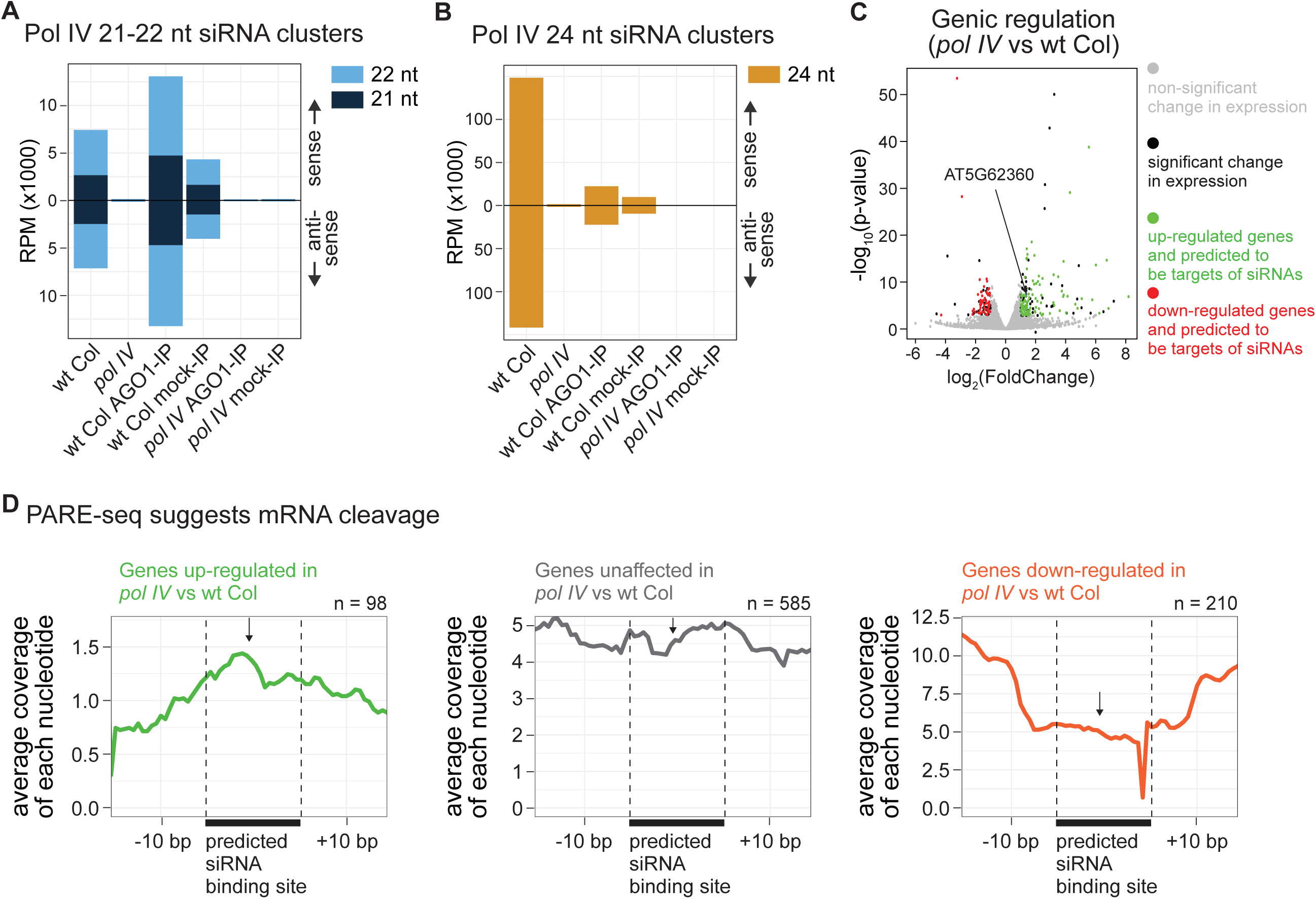
A potential role for Pol IV-dependent 21-22 nt siRNAs in post transcriptional gene silencing. A. AGO1 incorporation of Pol IV-dependent 21-22 nt siRNAs in wt Col and *pol IV* mutants. Mock IPs did not include a primary antibody. B. AGO1 incorporation of Pol IV-dependent 24 nt siRNAs in wt Col and *pol IV* mutants. C. Genic expression changes as evidenced by RNA-seq comparing wt Col and *pol IV* mutants. Black dots represent significant changes and gray dots represent non-significant changes. Green and red dots represent significantly up-regulated and down-regulated genes respectively for genes predicted to be targets of Pol IV siRNAs. D. Metaplot showing accumulation of PARE sequencing reads aligned to predicted target sites of Pol IV-dependent 21-22 nt siRNAs that show AGO1 incorporation. Genes identified as up-regulated (green), unaffected (grey) and down-regulated (red) in part C are aligned separately (left, center and right). The black arrow marks the predicted target cleavage site by Pol IV-dependent 21-22 nt siRNAs. The black bar at the bottom shows the predicted siRNA binding site used to align multiple genes. n denotes the number of transcripts analyzed towards the metaplot.

The second prediction states that for these Pol IV-derived 21-22 nt siRNAs, we could computationally identify target mRNAs and their predicted cleavage sites, although these programs identify false positives at a high rate. We identified 49,683 proposed target sites of the Pol IV-dependent 21-22 nt siRNAs that are enriched in AGO1 (from Figure 5A). This includes 23,668 distinct transcript models encompassing 18,167 total genes. We performed this analysis to inform our experiments below, however, the presence of small RNA target sites by itself provides no direct evidence of function.

The third prediction states that for at least a subset of the predicted target genes, their mRNA levels would increase in *pol IV* mutant plants, when the targeting siRNAs are not generated. We used publicly available mRNA-seq expression data [28] to identify genes that have increased steady-state transcripts in *pol IV* mutants compared to wt Col. Assuming that the steady-state transcript levels of the Pol IV siRNA target genes will increase in *pol IV* mutants, we overlapped the two sets of genes and found 117 genes with both a predicted target site for a Pol IV-dependent 21-22 nt siRNA, and the expected increase in transcript levels in the *pol IV* mutant (green, Figure 5C). We compared the fraction of genes with predicted target sites in the up-regulated set (green, Figure 5C) and unaffected set (grey, Figure 5C) and did not find an enrichment of target sites in up-regulated genes (data not shown). However, the increase in mRNA levels can be observed only for a subset of genes because of a) the false positives identified by the mRNA targeted prediction algorithm and b) AGO1-incorporated siRNAs could potentially cause translational repression instead of mRNA cleavage. Similar to the second prediction, expression data could not provide direct evidence of Pol IV-dependent mRNA degradation. Therefore, we aimed to further investigate the siRNA-mRNA interaction using parallel analysis of RNA end (PARE) sequencing (see below).

The fourth prediction states that for genes that increase in steady state mRNA abundance in *pol IV* mutants, their mRNA cleavage products could be detected specifically around the predicted siRNA target sites. To determine if the Pol IV-dependent 21-22 nt siRNAs could cause cleavage of the target mRNA akin to a Pol II-derived siRNA or microRNA in AGO1, we analyzed publicly available PARE mRNA cleavage data from wt Col inflorescence [29]. We defined three sets of genes to investigate, one with increased transcripts in *pol IV* (green, Figure 5C), a control with unaffected transcripts (grey, Figure 5C) and a second control with decreased transcript abundance in *pol IV* (red, Figure 5C). We aligned the target transcripts by the predicted Pol IV-dependent 21-22 nt siRNA target site and mapped the PARE sequences to these transcripts. We expected there to be increased signature of mRNA cleavage and thus PARE reads around the predicted cleavage site of the target genes. For the transcripts with the increased steady-state levels in *pol IV* mutants (green, Figure 5C), we indeed observed increased coverage of PARE sequences at the predicted siRNA binding site compared to flanking regions (green, Figure 5D). In contrast, we found no such change in coverage for the control gene set with unchanged transcript levels in *pol IV* (grey, Figure 5D) or decreased transcript levels in *pol IV* (red, Figure 5D). The combined data of AGO1 incorporation and cleavage transcripts at the predicted target site support a model that Pol IV-dependent 21-22 nt siRNAs may function in gene regulation.

## Discussion

We began our investigation of Pol IV-dependent 21-22 nt siRNAs based on a single publication that observed an anti-dogmatic siRNA accumulation pattern in pollen [15]. Our work has confirmed the Pol IV-dependence of many TE 21-22 nt siRNAs. These siRNAs differ from other TE 21-22 nt siRNAs that require Pol II transcription, such as in *ddm1* mutants [16]. In addition, we find that the Pol IV-dependent 21-22 nt siRNAs are direct products of unspliced Pol IV transcripts and are produced when TEs are transcriptionally silent (wt Col inflorescence) and transcriptionally activated (wt Col pollen). The 21-22 nt siRNAs are generated from the same regions with simultaneous overlapping production of 24 nt siRNAs, suggesting that the same Pol IV / RDR2 transcripts that are acted upon by DCL3 to generate 24 nt siRNAs are also processed by DCL2/DCL4 to generate 21-22 nt siRNAs.

A significant remaining question is why the size ratio of Pol IV-derived siRNAs is heavily skewed towards production of 21-22 nt siRNAs in pollen. Our analysis shows that the number of 21-22 nt siRNAs is not greatly increased in pollen, but rather the amount of 24 nt siRNAs is drastically reduced in pollen, skewing the ratio of 21-22 vs. 24 nt siRNAs (Figure 1B). The tissue-specific reduction of 24 nt siRNAs in pollen, while still retaining the Pol IV-dependent 21-22 nt siRNAs, is similar to a *dcl3* mutant in sporophytic tissue (Figure 3A-B), suggesting that a lack of DCL3 activity in pollen could result in the observed pattern. We observe a lack of DCL3 mRNA expression in pollen, however, several components required for the biogenesis of these siRNAs (such as the largest subunit of Pol IV) also fail to accumulate, necessitating future research. In addition, in pollen there is TE transcriptional activation [12,13], but the resulting Pol II transcripts are not responsible for generating the TE 21-22 nt siRNAs observed in pollen, and therefore the term *epigenetically-activated siRNA* (*easiRNA*) is not suitable for pollen siRNAs.

In sporophytic tissue, Pol IV transcripts generate high levels of 24 nt siRNAs and low levels of 21-22 nt siRNAs. We investigated the biological function of these Pol IV-derived 21-22 nt siRNAs, and found that in the absence of DCL3 and 24 nt siRNAs, 21-22nt siRNAs generated by Pol IV/RDR2/DCL2/DCL4 can participate in RdDM. Additionally, these 21-22 nt siRNAs may participate in the post-transcriptional regulation of genic mRNAs. We found evidence that these 21-22 nt siRNAs are loaded into AGO1 and at the predicted mRNA target sites increased transcript cleavage was detected. However, further research is needed to conclusively demonstrate that these 21-22 nt siRNAs can direct AGO1 to post-transcriptionally cleave genic mRNAs. Nonetheless, we conclude that these siRNAs may be RNAi potent, and can in theory target complementary invading TEs and quickly initiate an RNAi defense. A question remains regarding what would limit AGO1 activity and PTGS if Pol IV can generate gene-regulating 21-22 nt siRNAs? However, the levels of these siRNAs are only a fraction of known highly-abundant microRNAs and tasiRNAs. Therefore, even though mRNA cleavage can be detected, this may not be occurring on enough mRNA molecules to have a phenotypic consequence.

Although it is not understood why there is a shift in the size distribution of Pol IV siRNAs in pollen, the functional consequence of this shift is clear. Unlike 24 nt siRNAs, 21-22 nt siRNAs are incorporated into AGO1, which participates in gene regulation. Therefore, the function of Pol IV in pollen is likely to shift towards gene regulation (function of 21-22 nt siRNAs). Here we show evidence that Pol IV-derived 21-22 nt siRNAs may participate in post-transcriptional regulation. We speculate that this function may be connected to establishing hybridization barriers. Pol IV mutants fail to establish the triploid block, which ensures that the maternal : paternal ratio of genetic contribution is 2:1 in the early endosperm [15]. Pollen Pol IV-derived 21-22 nt siRNAs associate with gene expression changes [15], and TE siRNAs in pollen (of unknown biogenesis) are important regulators of imprinting through the post-transcriptional targeting of the genes UBP1b and PEG2 [30]. We propose that TE regions of the genome contribute towards diverse gene regulation via Pol IV-derived 21-22 nt siRNAs specifically in pollen and the early seed, including imprinting [31] and hybridization barriers [14,15].

## Methods

### Plants and materials

All plants used for small RNA sequencing as part of this study were grown in growth chambers with standard conditions: 22°C temperature and 16 hour light. Stage 1-12 inflorescence tissue was used for RNA isolation. *nrpd1a-3* (*pol IV*), *dcl1-9, dcl2-1, dcl3-1, dcl4-2, rdr1-1, rdr2-1, rdr6-15* alleles were used. Wt Col and *pol IV* have standard 1n pollen samples and 2n pollen generated using the *osd1* mutation [15], which were used as replicates in this study.

### AGO1 Immunoprecipitation

0.5g inflorescence tissue per sample was ground with liquid nitrogen and homogenized in lysis buffer (50mM Tris pH 7.5, 150mM NaCl, 5mM MgCl2, 10% glycerol, 1% IGEPAL, 0.5mM DTT, 1mM PMSF, and 1X GoldBio protease inhibitor) for 15 minutes. Lysates pre-cleared for 15 minutes with 50µl goat anti-rabbit magnetic beads (NEB). Pre-cleared lysates were then incubated with either goat anti-rabbit magnetic beads only (mock IP) or beads plus 5µg anti-AGO1 primary antibody (Agrisera)(AGO1 IP). IPs were performed at 4°C for 2 hours with end-over-end rotation. Beads were then washed three times 5 minutes in wash buffer (50mM Tris pH 7.5, 150mM NaCl, 5mM MgCl2, 0.5mM DTT). RNA was extracted directly from washed beads using TRIzol reagent, and small RNA libraries were constructed as described below directly from this RNA.

### Small RNA sequencing

Total RNA was extracted with phenol chloroform method using TRIzol reagent (Thermo Fisher Scientific). Small RNAs were enriched using miRVana miRNA isolation kit (Thermo Fisher Scientific). The TrueSeq Small RNA Library Preparation Kit (Illumina) was used to make sequencing libraries for total or IP-enriched small RNAs. Multiplexed libraries were sequenced on a HiSeq4000 (Illumina).

### Small RNA processing

Adapter TGGAATTCTCGGGTGCCAAGG was removed from demultiplexed libraries using fastx toolkit (http://hannonlab.cshl.edu/fastx_toolkit/). These sRNAs were mapped to the genome using bowtie 1.2.2 (-v 0) to determine the number of total genome matching reads which was used to normalize sRNA counts [32]. sRNA Workbench [33] was used to filter out low complexity reads, t/rRNA reads and retain 18-28 nt reads that match the Arabidopsis TAIR10 genome. ShortStack 3.8.5 [34] was used to map the sRNAs to the genome using the parameters --nohp --mmap f --bowtie_m all. Bowtie 1.2.2 was used by ShortStack. For the digital Northern in Figure 3C-D, size limit of 18-28 nt was not applied to allow the visualization of longer RNAs.

### Cluster Identification

ShortStack 3.8.5 was used to identify clusters of 24 nt and 21-22 nt sRNAs. All small RNA sequencing data used in this study were individually mapped to the Arabidopsis genome using ShortStack [34]. These mapped files were filtered to retain either only 24 nt reads or 21-22 nt reads. All the samples were then merged to create two merged mapped files, one each for 24 nt and 21-22 nt reads. These merged mapped files were used as input for ShortStack to identify clusters with the default parameters except for mincov (set to 10) and pad (set to 50). The identified clusters were then filtered for Arabidopsis miRNA loci from miRBase 22.1 [35]. The miRNA filtered cluster list was filtered for Pol IV-dependent clusters with the criteria that average accumulation of reads was at least 2-fold reduced in *pol IV* compared to wt Col.

### Whole genome DNA methylation analyses

We used inflorescence tissue to isolate DNA and perform MethylC-sequencing as previously described [43]. Statistics of the sequenced reads is shown in Supplementary Table 1. We identified differentially methylated regions (DMRs) using default parameters of methylpy program [44] available in github (https://github.com/yupenghe/methylpy). DMRs, were aligned by their edge and CHH methylation was calculated across the region in bins of 50 nt size and averaged across DMRs.

### RNA sequencing data analyses

mRNA sequencing data from GSE99691 [28] was reprocessed. Adapters were removed and the sequences were mapped to the genome using STAR 2.6.0c (parameters: -- outMultimapperOrder Random --outSAMtype BAM SortedByCoordinate -- outFilterMultimapNmax 50 --outFilterMatchNmin 30 --alignSJoverhangMin 3) [36]. Summarize Overlaps from GenomicFeatures [37] was used to count abundance of genic transcripts using annotation from JGI v11, Arabidopsis v167 TAIR10. DESeq2 [38] was used for differential expression analysis.

### Target prediction

A list of candidate sRNAs was prepared and used for target prediction. All 21-22 nt sRNAs enriched in wt Col IP samples (at least 5 raw counts in each of the two replicate of IP samples and >2-fold accumulation over mock IP) and from Pol IV dependent 21-22 nt clusters were labelled as candidate small RNAs. These siRNAs were further filtered for loss of accumulation in *pol IV* AGO1 IP samples (> 2-fold reduction in *pol IV* AGO1-IP). These 440 sRNAs were used is psRNA [39] for target prediction with default parameters.

### PARE data analyses

Raw data from GSM1263708 [29] was reprocessed. Adapter sequence was removed and reads with length less than 12 nt were discarded. The sequences were mapped to Arabidopsis transcripts (JGI v11, TAIR10, v167 transcripts) using bowtie 1.2.2 and the parameters -v 0, -a. Bedtools v2.25.0 [40] was used to count the accumulation of these degradome sequences on transcripts. This count was normalized by the total transcripts investigated in the gene set.

## Data Accessibility

All the sRNA sequenced generated for this study have been deposited to GEO with the accession number GSEXXXXXX. Publicly available small RNA-seq (GSE41755, GSE74398, GSE57191, GSE118705, GSE84122, GSE79780), RNA-seq (GSE99691), and PARE-seq (GSM1263708) were used.

## Acknowledgments

The authors thank Saima Shahid for her advice on bioinformatic approaches. This work was supported by grant MCB-1608392 to R.K.S from the U.S. National Science Foundation.

---

**Supplementary Figure 1:**
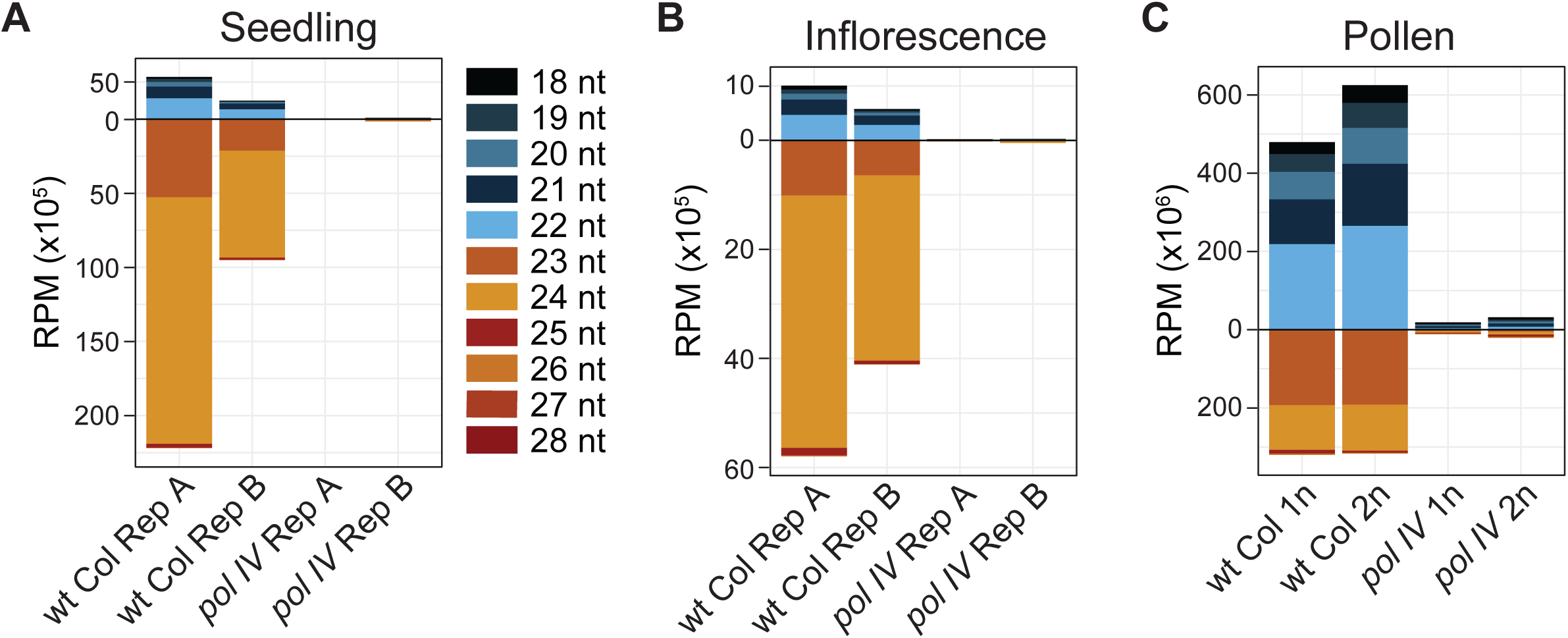
miRNA-normalized abundance of TE small RNAs. Reads per million (RPM) sequenced miRNAs of TE siRNAs in wt Col and *pol IV* mutant plants in three different tissue types - seedling, inflorescence and pollen. For each tissue type Pol IV-independent miRNAs were determined by comparing sRNA abundances between wt Col and *pol IV* mutant plants. The counts of Pol IV-independent miRNAs were then used to normalize tissue specific accumulation of TE siRNAs.

**Supplementary Figure 2:**
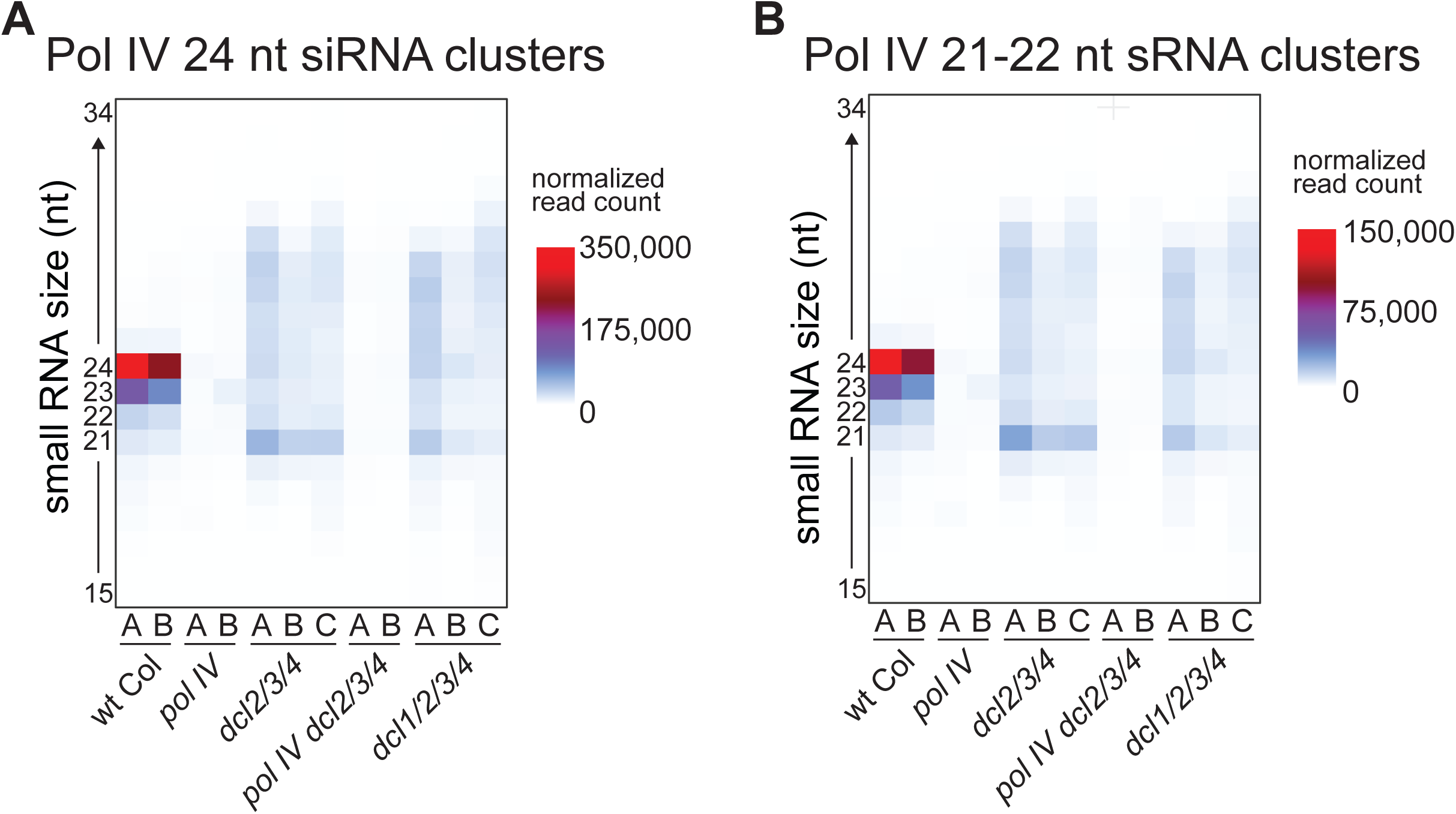
Investigation of Pol IV-dependent sRNAs in *dcl1/2/3/4* quadruple mutant. Heatmap of read counts of 15-34 nt small RNAs from Pol IV-dependent clusters in various dicer mutant combinations including the *dcl1/2/3/4* quadruple mutant from a seedling dataset (18). *dcl1/2/3/4* refers to the *dcl1/dcl2/dcl3/dcl4* quadruple mutant.

**Supplementary Figure 3:**
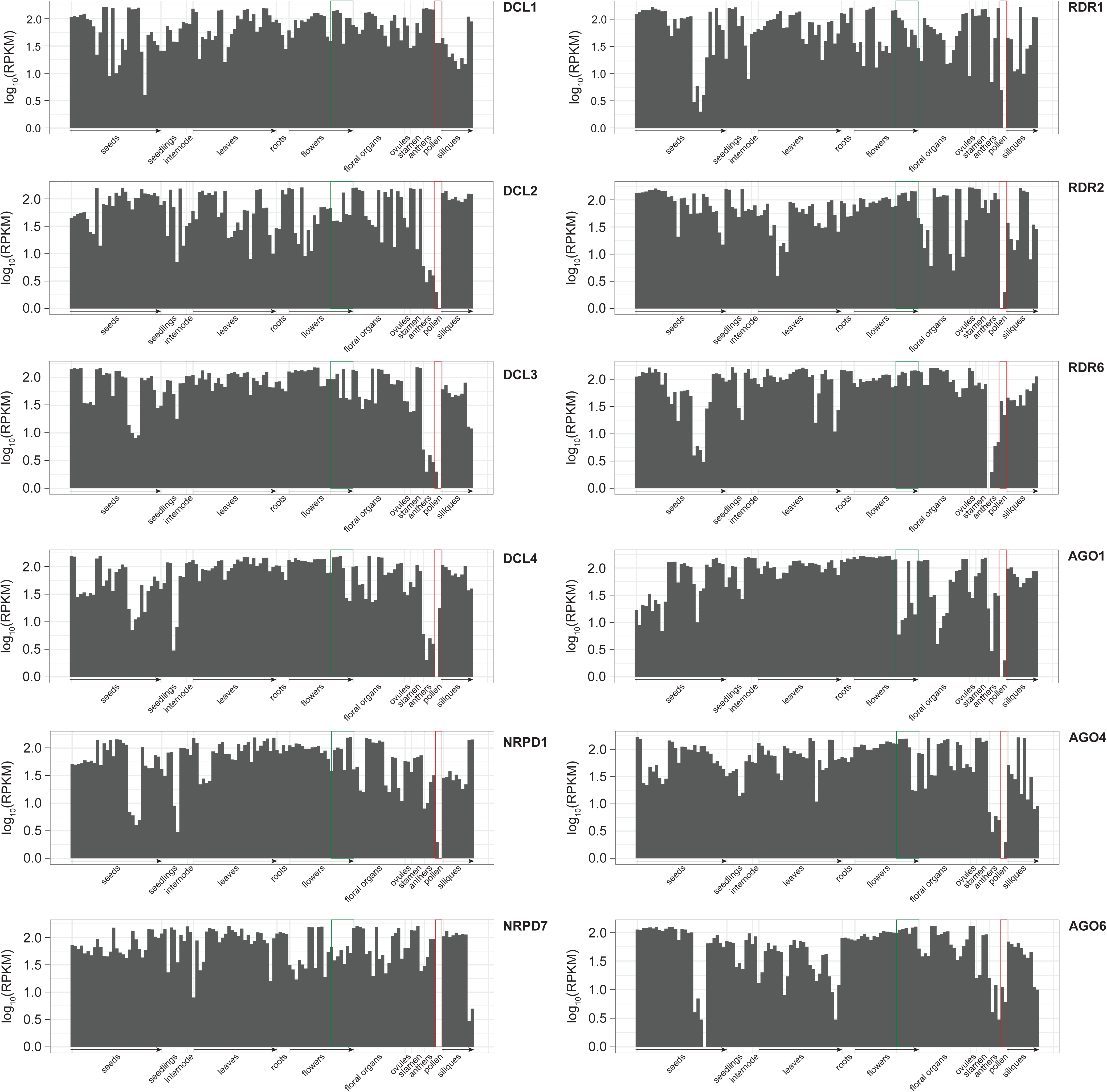
Developmental expression pattern of factors involved in Pol IV-dependent sRNA production. RPKM expression values of various proteins involved in Pol IV-dependent sRNA production of Pol IV-dependent RNA-directed DNA methylation (RdDM). Data were accessed from eFP-Seq Browser (42). Klepikova et al’s data (41) were used to show expression in differential developmental time points. Green box shows the inflorescence tissue used for almost all data reported in this study (unless specified otherwise). Red box highlights the pollen developmental time during which multiple factors are less expressed.

**Supplementary Figure 4:**
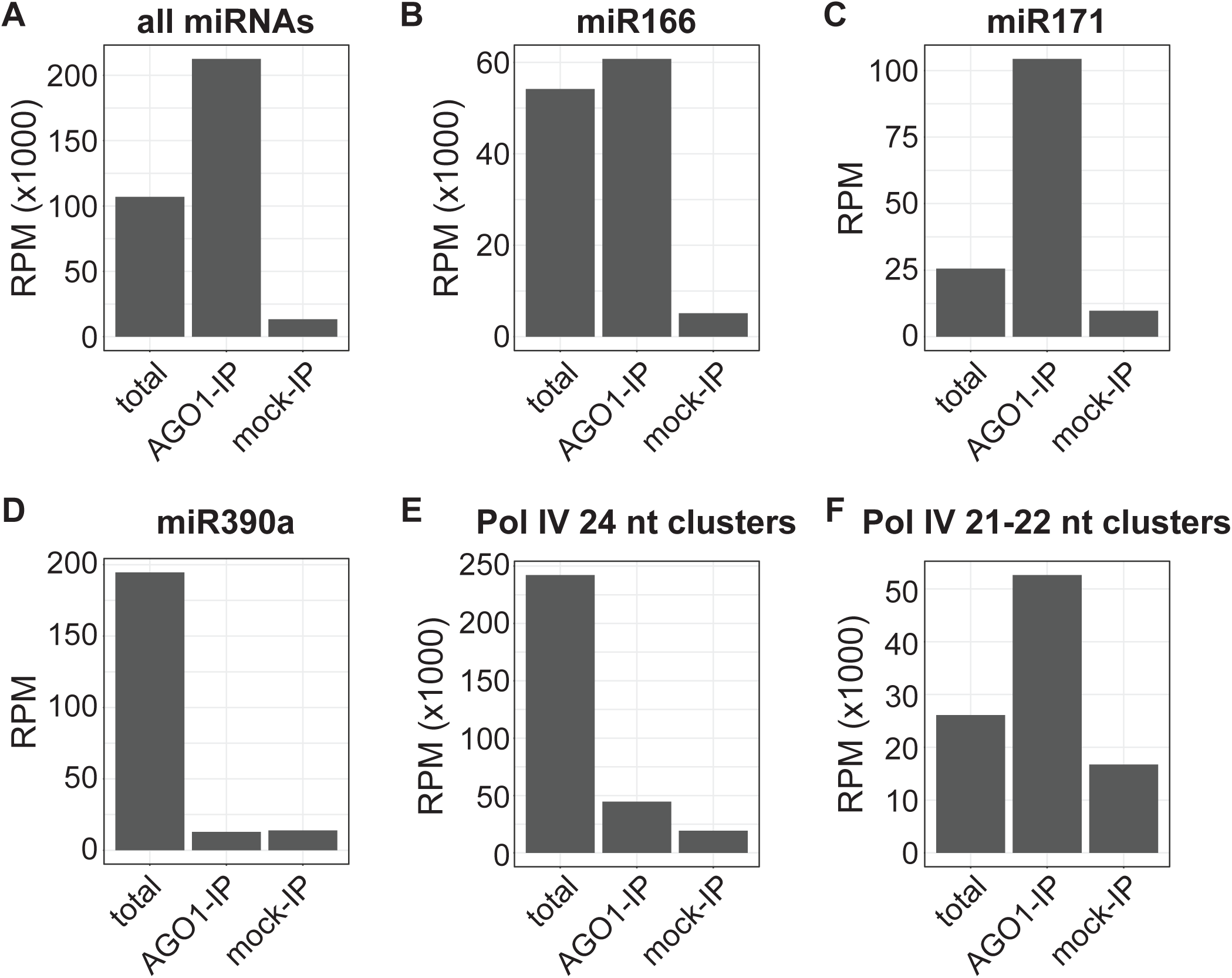
Pol IV 21-22 nt siRNAs are enriched in AGO1. sRNA accumulation from different loci is shown in total, AGO1-IP or negative control mock-IP samples in wt Col inflorescence tissue. A) High enrichment of all miRNAs in AGO1-IP vs mock-IP. B) and C) Positive controls miR166 and miR171 [45] show high enrichment of specific mature miRNA in AGO1. D) miRNA 390a is known to be incoporated specifically into AGO7 and not AGO1 [45,46] and is shown here as a negative control demonstrating no enrichment of mature miRNA in AGO1-IP vs mock. E) Only a subset of 24 nt sRNA from Pol IV 24 nt clusters are incorporated into AGO1-IP (AGO1-IP/total < 0.2) but the ones that do are enriched compared to mock (AGO1-IP-/mock IP > 2). and F) 21-22 nt sRNAs from Pol IV 21-22 nt clusters show genome-wide enrichment into AGO1 (AGO1-IP/total > 2) like all miRNAs in A. AGO1-IP/mock enrichment (>3) is not as high as miRNAs but higher than the negative control miR390a. Together these data demonstrate that Pol IV-dependent 21-22 nt sRNAs are incorporated into AGO1.

**Supplementary Table 1:** Sequencing statistics and availability.

| **Sample** | **Seq Approach** | **Figure(s)** | **Genome-**<br>**matched**<br>**reads** | **Genome**<br>**Coverage** | **Bisulfite**<br>**conversion**<br>**efficiency**<br>**(%)** | **Public**<br>**Availability** |
| --- | --- | --- | --- | --- | --- | --- |
| wt Col seedling Rep A | small RNA | 1A-B,<br>S2A-B | 20,569,377 | N/A | N/A | GSE74398 |
| wt Col seedling Rep B | small RNA | 1A-B,<br>S2A-B | 12,556,665 | N/A | N/A | GSE74398 |
| wt Col infl Rep A | small RNA | 1A-D, 3,<br>4B, 5A-B,<br>S4 | 4,856,770 | N/A | N/A | GSE118705 |
| wt Col infl Rep B | small RNA | 1A-D, 2B-F, 3, 4B.<br>5A-B, S4 | 6,627,617 | N/A | N/A | GSE118705 |
| wt Col pollen 1n | small RNA | 1A-B, 2B-F | 9,605,225 | N/A | N/A | GSE84122 |
| wt Col pollen 2n | small RNA | 1A-B | 18,627,821 | N/A | N/A | GSE84122 |
| pol IV seedling Rep A | small RNA | 1B, S2A-B | 20,609,065 | N/A | N/A | GSE74398 |
| pol IV seedling Rep B | small RNA | 1B, S2A-B | 10,528,909 | N/A | N/A | GSE74398 |
| pol IV pollen 1n | small RNA | 1B,<br>2B,C,E | 40,682,203 | N/A | N/A | GSE84122 |
| pol IV pollen 2n | small RNA | 1B | 38,768,834 | N/A | N/A | GSE84122 |
| wt Col infl Rep C | small RNA | 1C-D, 3,<br>4B, 5A-B | 5,793,758 | N/A | N/A | GSE118705 |
| wt Col infl Rep D | small RNA | 1C-D, 3,<br>5A-B | 5,686,327 | N/A | N/A | GSEXXXXXX |

**Supplementary Table 1: Sequencing statistics and availability**
| **Sample** | **Seq Approach** | **Figure(s)** | **Genome-**<br>**matched**<br>**reads** | **Genome**<br>**Coverage** | **Bisulfite**<br>**conversion**<br>**efficiency**<br>**(%)** | **Public**<br>**Availability** |
| --- | --- | --- | --- | --- | --- | --- |
| pol IV infl Rep A | small RNA | 1B-D,<br>2B,C,E, 3,<br>4B, 5A-B | 5,557,325 | N/A | N/A | GSE118705 |
| pol IV infl Rep B | small RNA | 1B-D, 3,<br>4B, 5A-B | 4,964,216 | N/A | N/A | GSEXXXXXX |
| ddm1 infl | small RNA | 2B-F | 5,823,347 | N/A | N/A | GSE79780 |
| ddm1/pol IV infl | small RNA | 2B-F | 10,119,843 | N/A | N/A | GSE79780 |
| dcl1 | small RNA | 3A-B | 8,529,042 | N/A | N/A | GSEXXXXXX |
| dcl2 | small RNA | 3A-B | 4,636,455 | N/A | N/A | GSE118705 |
| dcl3 | small RNA | 3A-B, 4B | 7,403,821 | N/A | N/A | GSE79780 |
| dcl4 | small RNA | 3A-B | 12,589,786 | N/A | N/A | GSEXXXXXX |
| dcl2/dcl4 | small RNA | 3A-B, 4B | 9,943,767 | N/A | N/A | GSEXXXXXX |
| dcl2/dcl3/dcl4 | small RNA | 3A-B, 4B | 3,505,470 | N/A | N/A | GSEXXXXXX |
| rdr1 | small RNA | 3C-D | 4,666,765 | N/A | N/A | GSEXXXXXX |
| rdr2 Rep A | small RNA | 3C-D<br>(including<br>heatmap) | 12,399,250 | N/A | N/A | GSE57215 |
| rdr2 Rep B | small RNA | 3C-D (not<br>in<br>heatmap) | 9,230,115 | N/A | N/A | GSE57215 |
| rdr6 Rep A | small RNA | 3C-D | 2,502,369 | N/A | N/A | GSE79780 |
| rdr6 Rep B | small RNA | 3C-D | 7,473,362 | N/A | N/A | GSE79780 |
| rdr1/rdr2/rdr6 Rep A | small RNA | 3C-D | 2,208,088 | N/A | N/A | GSEXXXXXX |
| rdr1/rdr2/rdr6 Rep B | small RNA | 3C-D | 2,847,374 | N/A | N/A | GSEXXXXXX |

**Supplementary Table 1: Sequencing statistics and availability**
| **Sample** | **Seq Approach** | **Figure(s)** | **Genome-**<br>**matched**<br>**reads** | **Genome**<br>**Coverage** | **Bisulfite**<br>**conversion**<br>**efficiency**<br>**(%)** | **Public**<br>**Availability** |
| --- | --- | --- | --- | --- | --- | --- |
| wt Col Rep A | methylC-seq | 4A | 19,989,099 | 25.13 | 99.74 | GSE118695 |
| wt Col Rep B | methylC-seq | 4A | 18,176,576 | 22.85 | 99.61 | GSE118695 |
| dcl2/dcl4 | methylC-seq | 4A | 19,886,103 | 25.00 | 99.59 | GSEXXXXXX |
| dcl3 Rep A | methylC-seq | 4A | 18,239,145 | 22.93 | 99.72 | GSE79746 |
| dcl3 Rep B | methylC-seq | 4A | 20,333,748 | 25.57 | 99.63 | GSEXXXXXX |
| dcl2/dcl3/dcl4 | methylC-seq | 4A | 15,837,317 | 19.90 | 99.86 | GSE118695 |
| pol IV | methylC-seq | 4A | 15,676,003 | 19.71 | 99.73 | GSE79746 |
| wt Col AGO1-IP Rep A | small RNA | 5A-B, S4 | 8,640,111 | N/A | N/A | GSEXXXXXX |
| wt Col AGO1-IP Rep B | small RNA | 5A-B, S4 | 5,020,914 | N/A | N/A | GSEXXXXXX |
| wt Col mock-IP Rep A | small RNA | 5A-B, S4 | 2,132,700 | N/A | N/A | GSEXXXXXX |
| wt Col mock-IP Rep B | small RNA | 5A-B, S4 | 4,338,460 | N/A | N/A | GSEXXXXXX |
| pol IV AGO1-IP Rep A | small RNA | 5A-B | 31,276,692 | N/A | N/A | GSEXXXXXX |
| pol IV AGO1-IP Rep B | small RNA | 5A-B | 8,214,904 | N/A | N/A | GSEXXXXXX |
| pol IV mock-IP Rep A | small RNA | 5A-B | 1,512,294 | N/A | N/A | GSEXXXXXX |
| pol IV mock-IP Rep B | small RNA | 5A-B | 2,297,768 | N/A | N/A | GSEXXXXXX |
| wt Col Rep A | RNA-seq | 5C | 13,035,982 | N/A | N/A | GSE99691 |
| wt Col Rep B | RNA-seq | 5C | 10,875,947 | N/A | N/A | GSE99691 |
| pol IV Rep A | RNA-seq | 5C | 10,999,691 | N/A | N/A | GSE99691 |
| pol IV Rep B | RNA-seq | 5C | 10,158,282 | N/A | N/A | GSE99691 |
| wt Col infl | PARE-Seq | 5D | 1,473,289 | N/A | N/A | GSE52342 |
| dcl2/dcl3/dcl4 seedling A | small RNA | S2A-B | 17,959,576 | N/A | N/A | GSE74398 |
| dcl2/dcl3/dcl4 seedling B | small RNA | S2A-B | 7,911,604 | N/A | N/A | GSE74398 |
| dcl2/dcl3/dcl4 seedling C | small RNA | S2A-B | 12,210,530 | N/A | N/A | GSE74398 |
| dcl1/dcl2/dcl3/dcl4 seedling A | small RNA | S2A-B | 9,771,195 | N/A | N/A | GSE74398 |
| dcl1/dcl2/dcl3/dcl4 seedling B | small RNA | S2A-B | 10,923,823 | N/A | N/A | GSE74398 |
| dcl1/dcl2/dcl3/dcl4 seedling C | small RNA | S2A-B | 11,359,174 | N/A | N/A | GSE74398 |

**Supplementary Table 1: Sequencing statistics and availability**
| **Sample** | **Seq Approach** | **Figure(s)** | **Genome-**<br>**matched**<br>**reads** | **Genome**<br>**Coverage** | **Bisulfite**<br>**conversion**<br>**efficiency**<br>**(%)** | **Public**<br>**Availability** |
| --- | --- | --- | --- | --- | --- | --- |
| pol IV/dcl2/dcl3/dcl4 seedling A | small RNA | S2A-B | 15,486,493 | N/A | N/A | GSE74398 |
| pol IV/dcl2/dcl3/dcl4 seedling B | small RNA | S2A-B | 11,736,662 | N/A | N/A | GSE74398 |

